# High delay discounting relates to core symptoms and to pulvinar atrophy in frontotemporal dementia

**DOI:** 10.1101/2024.05.16.594526

**Authors:** Valérie Godefroy, Anaïs Durand, Richard Levy, Bénédicte Batrancourt, Liane Schmidt, Leonie Koban, Hilke Plassmann

## Abstract

Behavioural variant frontotemporal dementia (bvFTD) is a neurodegenerative disorder characterized by behavioural changes and atrophy in brain regions important for decision-making. Computations such as trading off between larger later (LL) and smaller sooner (SS) rewards — called delay discounting in behavioural economics — might be heavily impaired by bvFTD. In this cross-sectional study, our objectives were to investigate (1) whether bvFTD patients show higher delay discounting than healthy controls, (2) whether this maladaptive discounting correlates with impulsivity-related bvFTD symptoms, and (3) in which brain regions atrophy is related to bvFTD’s steeper discounting. BvFTD patients (N=24) and matched controls (N=18) performed two delay discounting tasks: one with monetary rewards and one with food rewards. We compared discount rates (log(k)) in bvFTD patients and controls and tested their correlations with symptoms. We used participants’ structural MRI data and applied whole-brain mediation analyses to investigate brain structures mediating the effect of bvFTD on delay discounting. For both monetary and food rewards, delay discounting was significantly higher in bvFTD patients than in healthy controls. BvFTD patients’ higher discounting of both money and food was associated with their greater disinhibition and eating behaviour changes. Whole-brain mediation analyses revealed that (1) several brain regions (left thalamic pulvinar, left parahippocampal cortex, right temporal lobe) were predictive of steeper discounting of both money and food and (2) grey matter density in these brain regions, including most prominently the medial pulvinar, mediated the effect of bvFTD on discounting. The impulsive preference for sooner rewards captured by delay discounting might constitute a common mechanism of the behavioural symptoms of inhibition deficit and eating behaviour changes in bvFTD. Future studies could further investigate the potential role of medial pulvinar structural modifications as a transdiagnostic marker and a therapeutic target of impulsivity troubles.

## Introduction

Frontotemporal dementia (FTD) is the most common of a group of neurological conditions associated with predominant degeneration of the prefrontal and temporal regions.^1^ Behavioural variant FTD (bvFTD), the most common clinical variant, is characterized by significant changes in personality and behaviour. The main behavioural symptoms observed in bvFTD patients are disinhibition, apathy, loss of empathy, perseverative, stereotyped or compulsive behaviour, and eating behaviour changes.^2^ Behavioural changes in bvFTD patients often cause great distress to caregivers.^3,4^ Despite their highly negative impact on quality of life and caregiver well-being, safe and effective solutions to treat these symptoms are lacking.^5,6^ A more accurate understanding of the mechanisms underlying these symptoms could facilitate the development of more effective treatment strategies targeting bvFTD behavioural changes. The main goal of this paper is to explore the neuroanatomical correlates of impulsive decision-making as a core common factor of several behavioural symptoms of bvFTD.

The degree to which a patient displays impulsivity in decision-making can be approximated by measuring ‘delay discounting’, an approach from behavioural economics. Delay discounting refers to the extent to which people prefer smaller yet sooner over larger but later rewards.^7,8^ Individual differences in people’s tendency to discount delayed rewards can be captured by models from behavioural economics, such as the hyperbolic discounting model.^8–10^ The hyperbolic discounting model allows measurement of the discount rate k. Higher discount rates correspond to higher levels of impatience for reward or higher impulsivity. High discount rates have been detected in a number of psychiatric conditions, such as major depressive disorder, borderline personality disorder, bipolar disorder, bulimia nervosa and binge-eating disorder (see Amlung *et al*.^11^ for a meta-analysis). A growing body of literature has also suggested increased discount rates in neurodegenerative conditions such as Parkinson’s disease, Alzheimer’s disease and frontotemporal dementia (see Godefroy *et al*.^12^ for a review). As bvFTD patients tend to systematically choose immediate rewards rather than delayed rewards,^13^ they seem to constitute a good model for the study of high discounters.

How higher delay discount rates relate to bvFTD specific symptomatology and atrophy pattern is still unclear. Only a few studies have investigated delay discounting behaviour in bvFTD patients. Results suggest that bvFTD patients are steeper discounters and systematically more impulsive than healthy controls.^13–17^ These studies mostly focused on comparing patients with bvFTD to patients with Alzheimer’s disease (AD) to test the clinical value of using delay discounting to differentiate between these two conditions. Only one study used voxel-based morphometry to test correlations between discount rates and grey matter density in bvFTD and AD.^17^ This study found that AD patients showed increased discount rates compared to healthy controls under the influence of viewing emotionally negative pictures prior to choosing between SS and LL rewards. This increased impulsivity was associated with greater bilateral amygdala atrophy only in AD patients. In contrast, bvFTD patients discounted delayed rewards more than both AD patients and healthy controls, independent of the emotional context. However, this study did not aim to establish a specific pattern of brain structural changes leading to increased discount rates in bvFTD compared to healthy controls.

Against this background, in this paper we hypothesized that delay discounting is higher in bvFTD patients (compared to controls) and is correlated with at least two core behavioural symptoms of bvFTD: 1) disinhibition (or deficit of inhibition), which can be considered as a preference for the most immediate answers across various contexts (e.g., preference for immediate reactions of hostility and aggressiveness when confronted to frustration or irritation), and 2) eating behaviour changes such as binge eating and preference for sweet foods, which may correspond to a preference for immediately rewarding foods. We also predicted that the topography of brain damage in bvFTD patients could explain their higher discount rates. Using the statistical framework of whole-brain mediation analysis^18^ applied to structural MRI, we aimed at identifying the brain areas in which differences in structural topography between the patients and healthy controls explained group differences in discounting. The mediation framework can be very useful to uncover large-scale intermediate variables between a disease status and associated symptoms.^19^ Many behavioural symptoms, in particular disinhibition and impulsivity, are supposed to be closely associated with neurodegeneration in the orbitofrontal cortex (OFC) and ventromedial prefrontal cortex (vmPFC) in bvFTD.^20–22^ Building on prior work,^12^ we hypothesized that differences in discount rates between bvFTD patients and healthy controls are mediated by group differences in the neuroanatomy of these two brain regions in particular.

To test these hypotheses, we collected data capturing (1) discount rates for money and food rewards, (2) bvFTD behavioural symptoms, and (3) structural magnetic resonance imaging data (sMRI) from bvFTD patients and matched healthy controls. To compute discount rates, patients and healthy controls participated in two delay discounting tasks, with two different types of rewards (secondary monetary rewards and primary food rewards). We investigated reward impatience for two different rewards to investigate generalizability across reward domains. We estimated the discount rate k from each of these two tasks and compared them between patients and controls. We found that bvFTD patients were more impatient for both reward types. Next, we established a link between two core bvFTD behavioural symptoms related to impulsivity (i.e., lack of inhibition and eating behaviour changes) and discount rates. In a third step, we tested whether the differences in reward impatience between groups can be explained by changes in brain structure due to bvFTD.

## Materials and methods

### Participants

This study was part of a larger-scale protocol that also collected other data that are reported elsewhere.^21,23–25^ Participants were recruited in the context of a clinical study at the Paris Brain Institute in France (Clinicaltrials.gov: NCT03272230).^21,23–25^ This clinical study aimed at assessing several behavioural symptoms (disinhibition and apathy) and investigating their neural correlates in behavioural variant frontotemporal dementia. BvFTD patients were recruited in two tertiary referral centres in Paris: the Pitié-Salpêtrière Hospital and the Lariboisière Fernand-Widal Hospital. They were diagnosed according to the International Consensus Diagnostic Criteria.^2^ Inclusion criteria for bvFTD patients included presenting a Mini-Mental State Evaluation (MMSE) score of at least 20 to ensure that they would have the ability to undergo the full protocol. Healthy controls were recruited by public announcement; inclusion criteria were a MMSE score superior to 27 and matching the demographic characteristics of the bvFTD patient group (i.e., age, gender and education level). In total, 24 bvFTD patients (mean age = 66.6, 66.6% male) and 18 neurologically healthy controls (mean age = 62.6, 44.4% male) were recruited for the overall clinical study. Data were missing for some participants (total of four missing participants for each of the two intertemporal choice tasks) because of technical issues with the touch tablet used to collect the data of the intertemporal choice tasks. Hence, in this study, we used data from 22 bvFTD patients and 16 healthy controls for delay discounting of money and data from 21 bvFTD patients and 17 healthy controls for delay discounting of foods. (For additional details regarding the sample of 22 bvFTD patients and 17 healthy controls with at least one measure of delay discounting, refer to Supplementary Table 1.)

### Intertemporal choice tasks

Participants performed two intertemporal choice (ITC) tasks, one using monetary rewards and one using food rewards, in a randomized order. Both types of rewards were matched in economic value, meaning that one chocolate truffle used as a food reward had a retail price of about €1. Each of these tasks consisted of 32 choices between smaller sooner and larger later rewards (see Fig. 1). To incentivize participants to give us truthful answers, they were instructed that one of their 32 choices would be randomly selected at the end of the experiment, and the option they had chosen would be given to them at the indicated time. Thus, participants’ choices were incentive-compatible and non-hypothetical. During each choice, the two options were displayed for a maximum of 100 seconds. During this time, participants indicated their choice by pressing either a blue key on the keyboard with their right index finger to select the option on the left or a yellow key with their right middle finger to select the option on the right. Once the choice had been made, a message on the screen indicated the selected option. The maximum time between choice trials was either 3.5, 4 or 5s (randomized order) and depended on the participant’s response time for each choice. The values of the SS reward ranged from €8 to €35; the delay for the SS option was either 0, 14 or 28 days. LL options varied between €10 and €96; the delay for the LL option was either 14, 28 or 42 days. For the ITC task with food rewards, choice trial parameters were exactly the same except that reward amounts in euros were replaced by numbers of chocolate truffles with the economic value of €1 each. Supplementary Table 2 details all combinations of SS and LL options and delays that constituted the choice trials of the task, as well as the corresponding indifference k’s (k’s for which the presented LL and SS options would be chosen with equal probability). For each of the two ITC tasks, choice trials were presented in randomized order.

**Figure 1.**
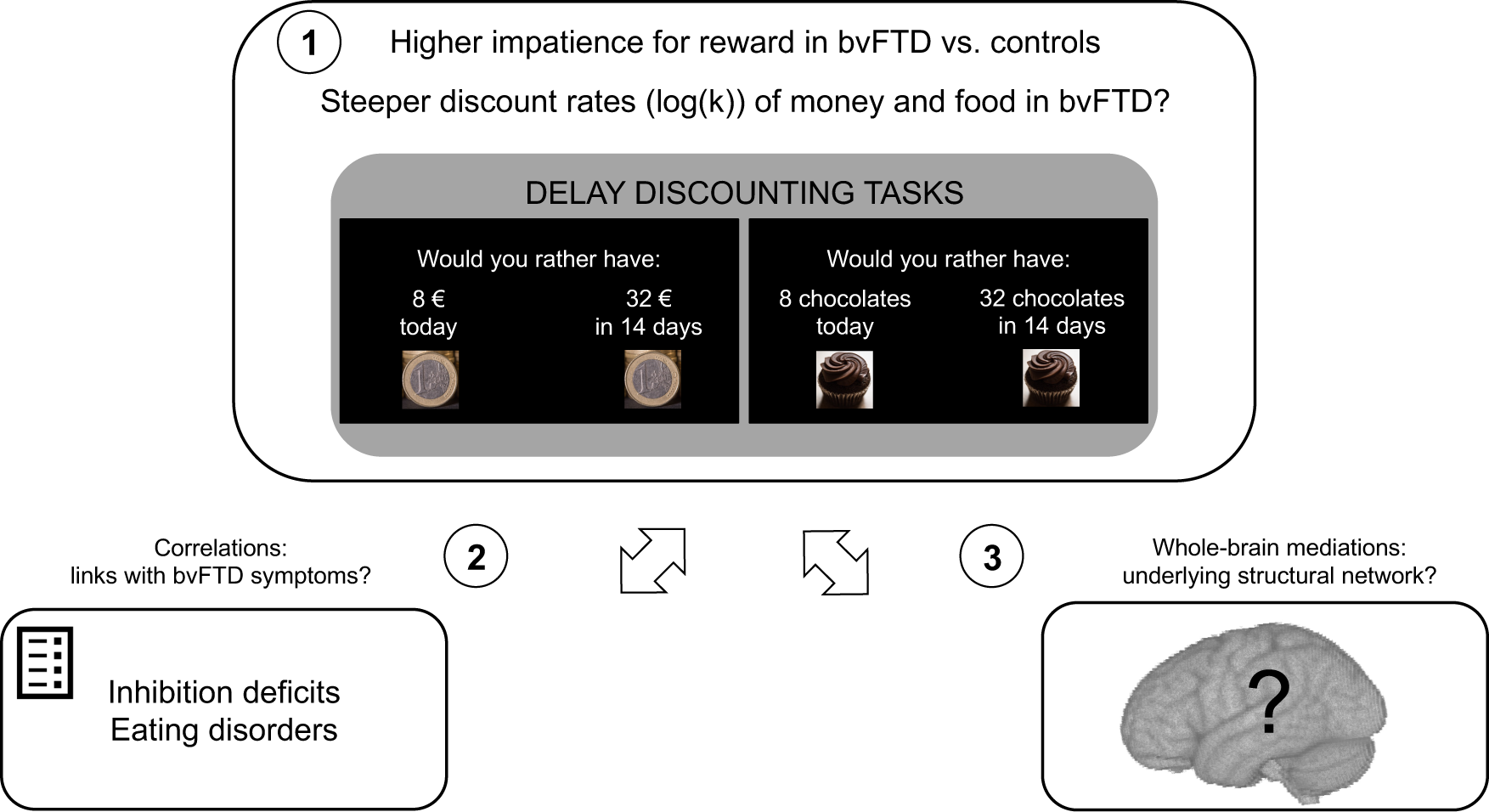
Study objectives and paradigm description. Study objectives are numbered from 1 to 3. Objective #1 was to examine the effect of bvFTD on discount rates for money and food rewards. The visual presentation of the two ITC tasks used to assess delay discounting in the study shows an example of trial with monetary rewards on the left and with food rewards (chocolate) on the right. Objective #2 was to show the links between higher delay discounting and core bvFTD behavioural symptoms suggestive of impulsivity: 1) lack of inhibition (measured by Hayling test error score) as a tendency to prefer immediate prepotent answers and 2) eating behaviour changes as a tendency to prefer immediate food rewards. Objective #3 was to investigate the brain regions in which atrophy may cause the increase in discount rates in bvFTD patients (compared to controls). For this purpose, we used a whole-brain mediation approach applied to participants’ grey matter density maps (T1 pre-processed for VBM).

### Measures of impulsivity-related bvFTD symptoms

We used measures of two behavioural symptoms of bvFTD^2^ that we assumed to be associated with impatience in decision-making: (1) the deficit of inhibition measured by the Hayling Sentence Completion Test (HSCT) and (2) eating behaviour changes measured by the Eating Behavior Inventory (EBI).

The Hayling Sentence Completion Test^26^ asks participants to complete 15 sentences using the appropriate word, as fast as possible (automatic condition, part A — e.g., for ‘The rich child attended a public ’, the correct answer is ‘school’), and 15 sentences using a completely unrelated word (inhibition condition, part B — e.g., for ‘London is a very lively —’, ‘city’ is considered an incorrect answer, but ‘banana’ would be considered correct). We selected the Hayling error score (number of errors in part B) as a measure of the difficulty in inhibiting a prepotent immediate response, as in Flanagan *et al*.^27^. Among the different tests available to assess inhibition deficits, the Hayling test presents several advantages: it is an objective measure, and impaired performances are distinctive characteristics of bvFTD patients.^28^

The Eating Behavior Inventory is a questionnaire with 32 questions investigating recent changes in four domains of eating behaviour: eating habits (e.g., ‘Seeks out food between meals’), food preference (e.g., ‘Is more attracted by sweet foods’), table manners (e.g., ‘Is eager to start eating’), and food approach (e.g., ‘Eats closer to his plate’).^29^ This questionnaire is particularly adapted to detect the specific eating behaviour changes of bvFTD and has been evidenced as a tool helping the differential diagnosis of this condition. For bvFTD patients, this questionnaire was completed by the caregiver to avoid self-report biases due to anosognosia in patients.

In order to investigate the specificity of the link between discount rates and these two impulsivity-related behavioural symptoms, we also tested the links between discount rates and apathy, another core behavioural symptom of bvFTD. As apathy is known to be a complex multidimensional symptom, we used the Dimensional Apathy Scale (DAS)^30^ to measure three subtypes of apathy derived from the theoretical model proposed by Levy and Dubois^31^: initiation apathy (deficit of initiation of thoughts and actions; 8 items; e.g., ‘I set goals for myself’ as reversed item), emotional apathy (emotional blunting; 8 items; e.g., ‘I become emotional easily when watching something happy or sad on TV’ as reversed item), and executive apathy (impairment in executive functions to manage goals; 8 items; e.g., ‘I find it difficult to keep my mind on things’).

### MRI data acquisition and pre-processing

Brain imaging data were acquired at the neuroimaging centre (CENIR) of the Paris Brain Institute with a Siemens Prisma whole-body 3T scanner (with a 12-channel head coil). Structural images were acquired using a T1 weighted MPRAGE sequence (TR 2400 ms; TE 2.17 ms; FOV 224 mm; 256 slices; slice thickness 0.70 mm; TI 1000 ms; flip angle 8°; voxel size 0.7 mm isomorphic; total acquisition time 7:38). T1 images were pre-processed for voxel-based morphometry (VBM) analyses using the Statistical Parametric Mapping (SPM) software (version 12). The pre-processing consisted of several standardized steps. Image files were first segmented and registered using rigid linear deformations. These images were then used as input to create a customized Dartel template and individual flow fields for each subject. The Dartel module determines the nonlinear deformations for warping all the grey and white matter images so that they match each other. Finally, spatially normalized and smoothed Jacobian scaled grey matter images were generated, using the deformations estimated in the previous step.

### Data analyses

The data analyses were performed in several steps (see Fig. 1). First, we verified that bvFTD patients were more impatient for smaller sooner rewards as compared to controls. Second, we tested whether bvFTD symptoms suggestive of impulsivity (based on the Hayling Sentence Completion Test and the Eating Behavior Inventory) and delay discounting were positively correlated. Third, we investigated in which brain regions the grey matter atrophy mediated the effect of bvFTD on preference for sooner options. All the analyses were performed using R Studio (1.2.1335) and Matlab (R2017b) toolboxes.

### Computation of discount rates

For each participant and for each reward type we first computed his or her discount rate k. For each choice, we computed the corresponding theoretical ‘indifference’ discount rate k (i.e., the theoretical k value for which an individual would consider both options of the trial as equivalent assuming a hyperbolic discount function where *V = A* / (1 *+ kD*)) using the following equation:

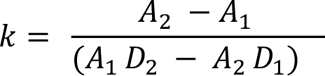

where *A_1_* = absolute value of SS reward, *A_2_* = absolute value of LL reward, *D_1_* = SS delay and *D_2_* = LL delay. We fitted a logistic probability function relating the theoretical indifference k value of a trial to the probability of choosing the LL option. We then used this function in each participant to identify their indifference point, that is, the theoretical indifference k value at which their probability of LL choice was equal to 50% (and thus equal to the probability of SS choice). For some participants showing patterns of unique answers (i.e., only SS or only LL choices) or very inconsistent answers, we could not use this method of estimation. We took advantage of the strong linear relationship between %SS choice and k, and within each population sample (bvFTD and controls) we imputed k values where they were missing. For participants with 0% SS choice, the imputed value was the minimum k observed in the rest of the sample; for participants with 100% SS choice, the imputed value was the maximum k. For participants with very inconsistent answers, we used a linear interpolation method to approximate missing k’s from %SS values. Because k values are generally not normally distributed, we used the log-transformed values of k (log(k)).

For both the money and food paradigms, we verified that the computed discount rates were related to how sensitive they were to LL reward (see Supplementary Methods). To compute sensitivity to LL reward, we fitted a logistic regression model in each participant predicting the trial-to-trial probability of choosing the LL option from the LL reward amount and LL delay value. We used the regression coefficient of LL reward amount as an estimate of the individual sensitivity to LL reward (higher sensitivity to reward corresponding to higher values).

### Effects of bvFTD on discount rates

Our hypothesis was that higher discount rates would be observed in bvFTD patients compared to controls. We compared discount rates in bvFTD patients and controls using non-parametric Wilcoxon tests.

### Symptom correlation with increased discount rates in bvFTD

Our hypothesis was that bvFTD symptoms of inhibition deficits and eating behaviour changes were related to higher discount rates in bvFTD. We used non-parametric Spearman rank correlations to test the links between the participants’ discount rates (log(k)) for money and for food rewards and (1) the Hayling error score (measuring inhibition deficit) and (2) the EBI total score (measuring eating behaviour changes). We tested these associations across both groups (bvFTD and controls) and within each group. We also used non-parametric Spearman rank correlations to test the links between the participants’ discount rates and apathy subtypes measured by DAS subscales. We predicted that discount rates would significantly correlate only with the executive apathy subtype (and not with the initiation and emotional apathy subtypes), as it is conceptually related to impulsivity as a measure of executive dysfunctions (e.g., the difficulty to inhibit prepotent responses).

### Structural brain mediators of increased discount rates in bvFTD

To investigate which brain regions contribute to the differences in discount rates between bvFTD patients and healthy controls, we used a whole-brain mediation analysis^18^ applied to structural MRI data. In this approach, each brain voxel is tested as a potential mediator of the effect of bvFTD on discount rates. This approach allowed us to identify regions across the whole brain in which grey matter density (GMD) loss explained the effect of bvFTD condition on discount rates (i.e., path ab, mediation effect). This analysis also allowed us to identify brain regions in which GMD would be different between the groups (i.e., path a, group effect) and predictive of the discount rate controlling for group effect (i.e., path b, intrinsic link between neuroanatomy and delay discounting). We were interested in brain structures being involved in all three effects.

To this end, participants’ GMD maps (T1 images pre-processed for VBM) were submitted to a single-level whole-brain mediation analysis. We used a bootstrapping procedure (with 10,000 generated samples) to detect the most robust mediators. Resulting statistical maps were thresholded at p < 0.05 FDR-corrected across the whole brain and across path a, b and ab analyses (corresponding to a voxel level of p < 0.008 for both rewards). To conduct the mediation analyses, we used the CANlab Mediation toolbox available at https://github.com/canlab.

## Results

### Higher discount rates in bvFTD patients than in controls

Participants’ choices allowed us to estimate their discount rate (log(k)). We observed a frequent ‘mono-choice’ pattern of answers (only SS or only LL options chosen throughout the 32 trials); seven bvFTD patients exclusively chose the SS option for the monetary task, and five bvFTD patients exclusively chose the SS option for the food task. Moreover, inconsistent patterns of answers for which the hyperbolic discount model did not fit (e.g., with the probability of choosing the LL option decreasing with larger amounts of LL rewards) were observed only in bvFTD patients (five for the money paradigm and six for the food paradigm).

For monetary rewards, bvFTD participants had a mean log(k) parameter of -2.25 (median log(k)=-2.11, corresponding to a k of 0.12, range=[-3.35, -1.73]), and healthy controls showed a mean log(k) parameter of -3.33 (median log(k)=-3.05, corresponding to a k of 0.05, range=[-6.58, -1.04]). For food rewards, bvFTD participants had a mean log(k) parameter of -1.63 (median log(k)=-1.54, corresponding to a k of 0.21, range=[-2.60, -1.16]), and healthy controls showed a mean log(k) parameter of -3.00 (median log(k)=-2.33, corresponding to a k of 0.10, range=[-7.49, -1.16]).

We verified that, for the money paradigm, increased discount rate was associated with lower sensitivity to LL reward (R=-0.59, p=0.0001, 95% CI=[-0.77, -0.35]), even after correcting for group effect (R=-0.34, p=0.04, 95% CI=[-0.66, -0.005]) (see Supplementary Fig. 1). For the food paradigm, after removing the extreme outliers (two on discount rate and three on sensitivity to LL reward), increased discount rate was also associated with lower sensitivity to LL reward (R=-0.75, p=5.10^-7^, 95% CI=[-0.88, -0.54]), including after correcting for group effect (R=-0.55, p=0.001, 95% CI=[-0.81, -0.18]) (see Supplementary Fig. 1). These significant links with sensitivities to LL reward confirm that the computed discount rates are representative of the tendency to prefer most immediate options regardless of the offered LL rewards.

BvFTD patients had higher discount rates and thus higher impatience for reward compared to controls for both money rewards (W=243, p=0.048, effect size=0.32) (see Fig. 2.A) and food rewards (W=265, p=0.01, effect size=0.42) (see Fig. 2.B). Relatedly, bvFTD patients showed lower sensitivity to LL rewards than controls for both money and food rewards (see Supplementary Fig. 2). Moreover, we performed two robustness analyses for the comparison of bvFTD patients and controls: 1) As bvFTD and control groups with available ITC data were not matched on age, we checked that differences in discount rates between bvFTD patients and controls were still significant while controlling for the effect of age (B=0.95, p=0.03, 95% CI=[0.08, 1.81] for money; B=1.29, p=0.008, 95% CI=[0.35, 2.23] for food), and 2) We checked that the difference in discount rate for food between bvFTD patients and controls was still significant even after removing two extreme outliers (with very low discount rates) among controls (W=223, p=0.04, effect size=0.35).

**Figure 2.**
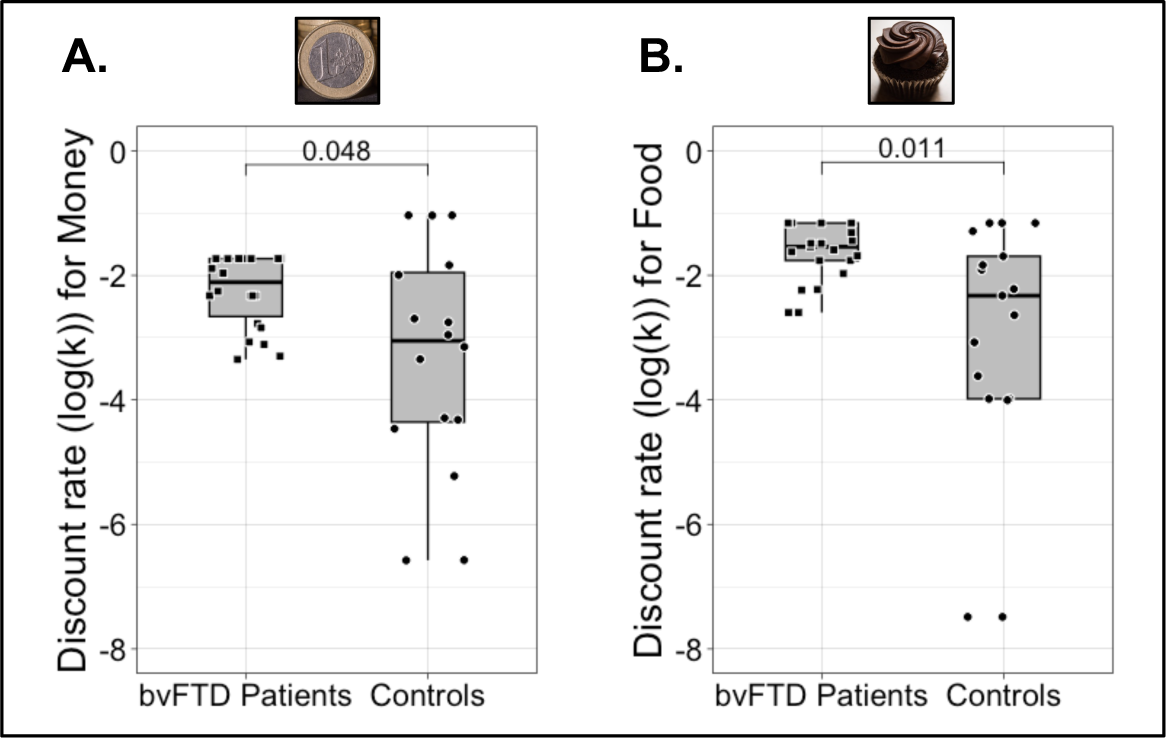
Effect of bvFTD on discount rates (log(k)). **A)** Wilcoxon test of the difference between bvFTD patients (N=22) and controls (N=16) on log(k) with money rewards; log(k)- Money was higher in bvFTD patients than in controls (W=243, p=0.048, effect size=0.32). **B)** Wilcoxon test of the difference between bvFTD patients (N=21) and controls (N=17) on log(k) with food rewards; log(k)-Food was higher in bvFTD patients than in controls (W=265, p=0.01, effect size=0.42). For each box plot, the lowest horizontal line represents the first quartile (Q1), the middle horizontal line represents the median, and the highest horizontal line is the third quartile (Q3). The lowest end of the box plot is defined as max(min, Q1-1.5*(Q3-Q1)); the highest end corresponds to min(max, Q3+1.5*(Q3-Q1)). Black dots correspond to individuals.

### Lack of inhibition and eating behaviour changes correlate with bvFTD patients’ increased discount rates

Participants showing higher deficits of inhibition showed higher discount rates for both rewards (money: R=0.54, p=0.0006, 95% CI=[0.25, 0.74]; food: R=0.39, p=0.02, 95% CI=[0.04, 0.67]). Participants with more eating behaviour changes also showed higher discount rates for both rewards (Money: R=0.38, p=0.02, 95% CI=[0.01, 0.67]; Food: R=0.46, p=0.003, 95% CI=[0.13, 0.73]) (see Fig. 3 from A to D). Of note, correlations between discount rates for food and both symptoms were still significant even after removing two extreme outliers (two controls) on the food discount rate (with deficits of inhibition: p=0.04; with eating behaviour changes: p=0.02). Correlations with the largest effect sizes (still significant after Bonferroni correction for multiple testing) were between inhibition deficits and discount rates for money as well as between eating behaviour changes and discount rates for food rewards. For these two correlations, we further tested the links among bvFTD patients and among controls separately (see Supplementary Fig. 3). We found that among bvFTD patients higher lack of inhibition was correlated with higher discount rates for money (R=0.67, p=0.0009, 95% CI=[0.36, 0.85]). The discount rates for food were not correlated with eating behaviour changes within the group of bvFTD patients but they were within healthy controls (R=0.52, p=0.03, 95% CI=[0.006, 0.86]).

**Figure 3.**
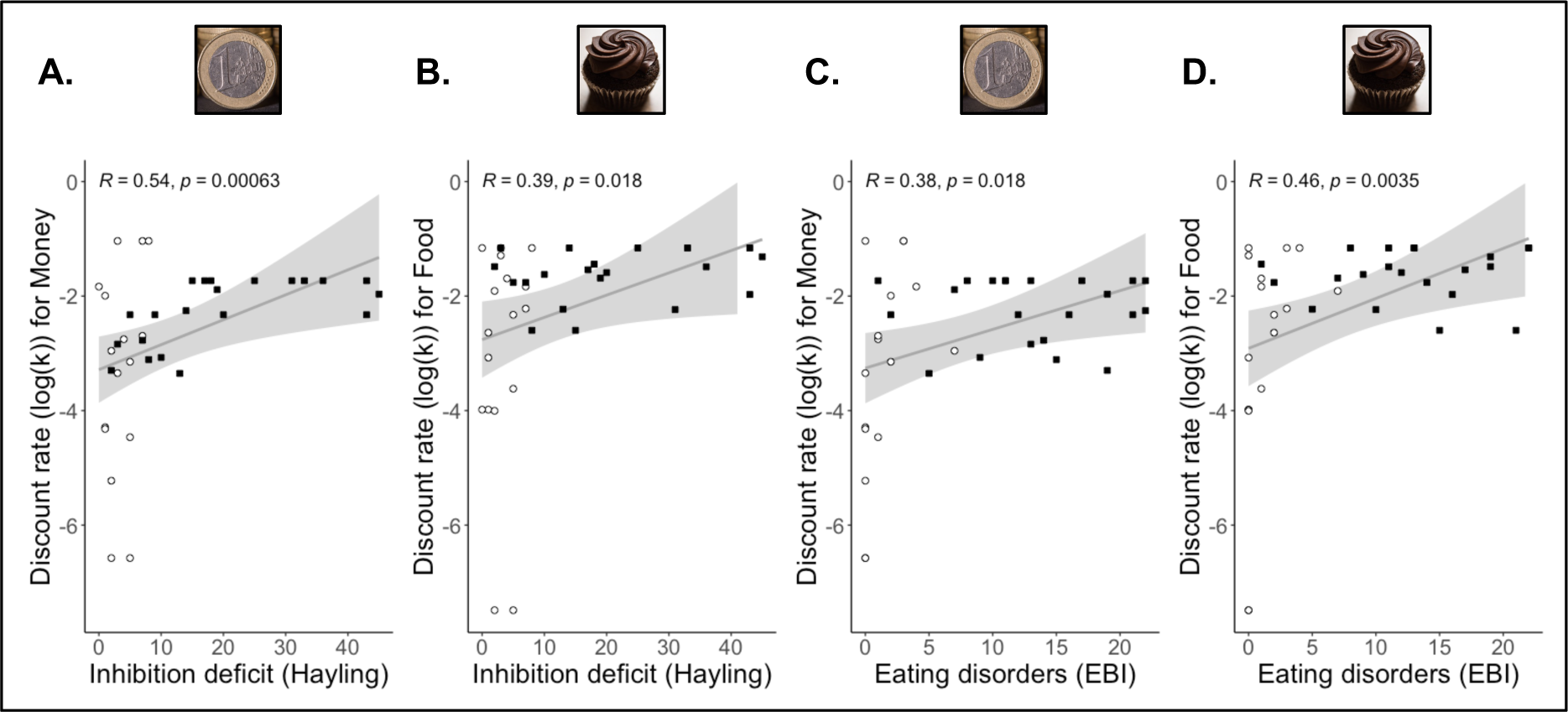
Relationship between bvFTD impulsivity symptoms and discount rates (log(k)). **A)** Spearman correlation between inhibition deficit (measured by Hayling error score) and discount rate for money across bvFTD patients (N=22; represented as black squares) and controls (N=16; represented as white circles) (R=0.54, p<0.001, 95% CI=[0.25, 0.74]); higher inhibition deficit (or preference for immediate prepotent answers) was related to higher log(k)- Money. **B)** Spearman correlation between inhibition deficit (measured by Hayling error score) and discount rates for food across bvFTD patients (N=21; represented as black squares) and controls (N=17; represented as white circles) (R=0.39, p=0.02, 95% CI=[0.04, 0.67]); higher inhibition deficit (or preference for immediate prepotent answers) was related to higher discount rates for food. **C)** Spearman correlation between eating behaviour changes (measured by EBI questionnaire) and discount rates for money across bvFTD patients (N=22; represented as black squares) and controls (N=16; represented as white circles) (R=0.38, p=0.02, 95% CI=[0.01, 0.67]); participants with more eating behaviour changes (or preference for immediate food rewards) had higher discount rates for money. **D)** Spearman correlation between eating behaviour changes (measured by EBI questionnaire) and discount rates for foods across bvFTD patients (N=21; represented as black squares) and controls (N=17; represented as white circles) (R=0.46, p=0.003, 95% CI=[0.13, 0.73]); participants with more eating behaviour changes (or preference for immediate food rewards) showed higher discount rates for foods.

To better disentangle the links between reward impatience and eating behaviour changes, we further explored the correlations between the four subscales of EBI and discount rates for food (see Supplementary Fig. 4). The *eating habits* (R=0.42; p=0.008; 95% CI=[0.09, 0.70]), *table manners* (R=0.40; p=0.01; 95% CI=[0.05, 0.65]) and *food approach* (R=0.49; p=0.002; 95% CI=[0.18, 0.72]) subscales were significantly correlated with discount rates for food, but the *food preference* subscale was not (R=0.28; p=0.09; 95% CI=[-0.07, 0.58]). Thus, it is mostly the disinhibited aspect of eating behaviour changes (or the preference for immediate ingestion of food independent of its taste) that seems to be linked to reward impatience.

Finally, we investigated correlations between discount rates and measures of apathy subtypes (see results in Supplementary Fig. 5). As expected, discount rates of both money and food rewards were significantly associated only with the *executive* apathy subtype (i.e., deficit of executive functions). They were not significantly related to *initiation* apathy (i.e., deficit of self-initiation of actions and thoughts) or to *emotional* apathy (i.e., emotional blunting). This suggests that discount rates are more closely linked with impulsivity-related symptoms than with other symptoms in which impulsivity is not supposed to be involved.

### Brain regions contributing to bvFTD patients’ increased discount rates

We used whole-brain mediation analysis to test which brain regions’ GMD mediated the effect of bvFTD (compared to controls) on the discount rate (log(k)) for both the money and food rewards (see Methods). We identified the brain regions involved in three statistical paths (FDR-corrected, q<0.05, across paths a, b and ab) (see Fig. 4 and see Fig. 5). As expected, path a, which characterizes the effect of bvFTD on brain GMD, showed a very broad pattern of atrophy in bvFTD, including frontal regions (OFC, vmPFC, anterior cingulate cortex, dorsomedial PFC, dorsolateral and lateral PFC, insula), temporal regions (mostly anterior temporal) and subcortical regions (striatum, amygdalae, thalamus, hippocampus). Path b shows the pattern of regions in which GMD predicts the discount rate, controlling for the effect of group. For path b, we were mostly interested in negative contributors (i.e., lower GMD in these regions was related to higher discount rates) as they were potential positive mediators of bvFTD patients’ increase in discount rates (resulting from the conjunction of negative path a and negative path b). Finally, path ab represents brain GMD, mediating the effect of bvFTD condition on discount rate. Most regions found to be mediators were positive mediators (i.e., they positively contribute to the increase in discount rates resulting from the effect of bvFTD condition on GMD). Clusters identified for paths b and ab for money and food stimuli are further detailed in Supplementary Tables 3 and 4.

**Figure 4.**
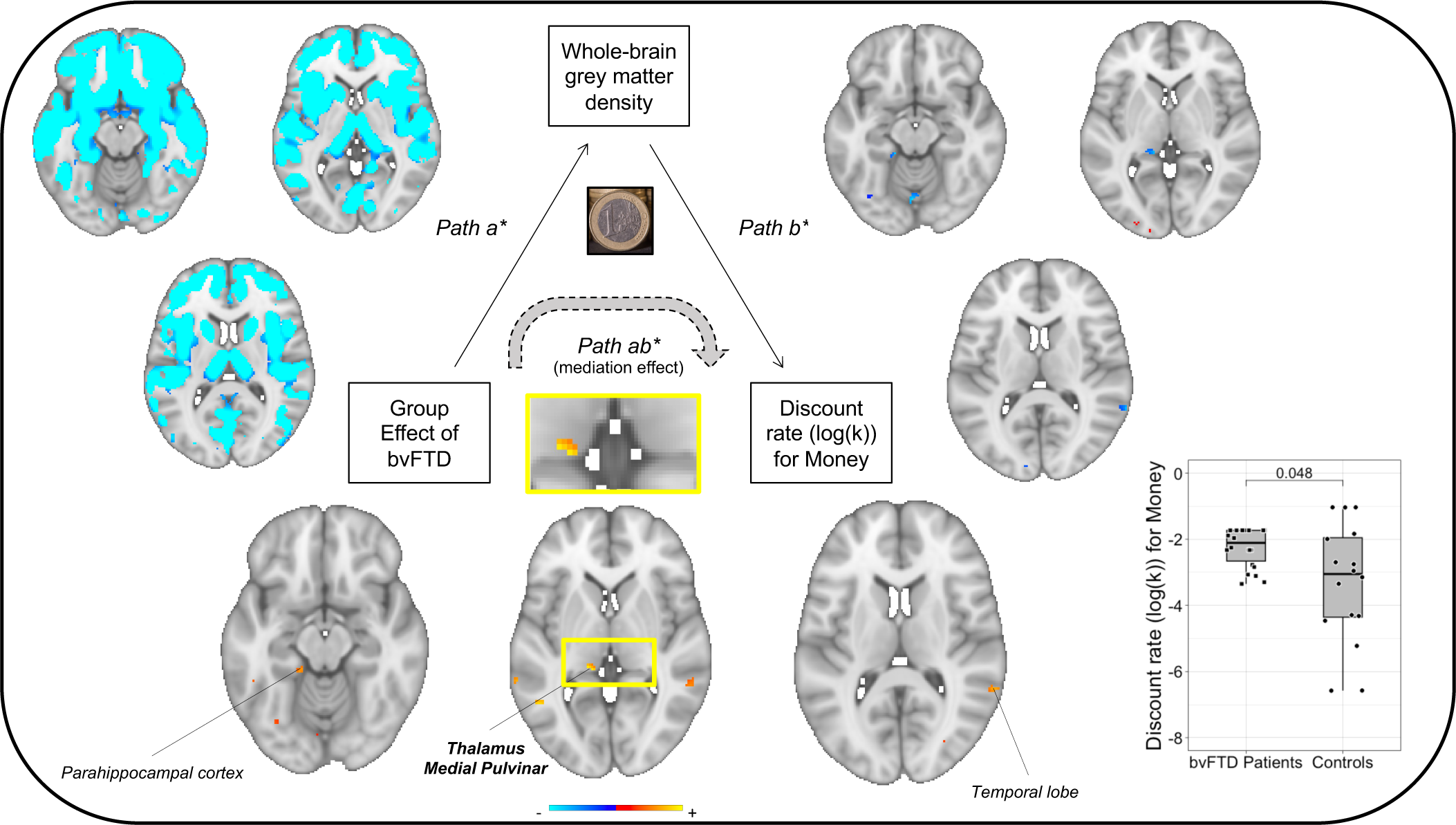
Neuroanatomical whole-brain mediators for the effect of bvFTD on discounting of monetary rewards. The path diagram displays the voxels of grey matter density that significantly contributed to each path of the mediation model. The blue voxels in *path a* reflect the decreased GMD in bvFTD (n=22) compared to healthy controls (n=16), which corresponds to the typical atrophy pattern observed in bvFTD. *Path b* shows the voxels in which GMD is correlated with the discount rates across all participants (n=38) by controlling for the effect of group. Voxels in blue correspond to a negative correlation in which lower grey matter density contributes to higher discounting. *Path ab* represents brain regions in which GMD mediates the effect of bvFTD on discount rates. Voxels in yellow correspond to positive mediators that positively contribute to the increase in discounting due to bvFTD. *All results are FDR- corrected and thresholded at q<0.05 across the three paths. The graph on the left shows the significant direct effect of bvFTD condition (compared to healthy controls) on the discount rate of money rewards.

**Figure 5.**
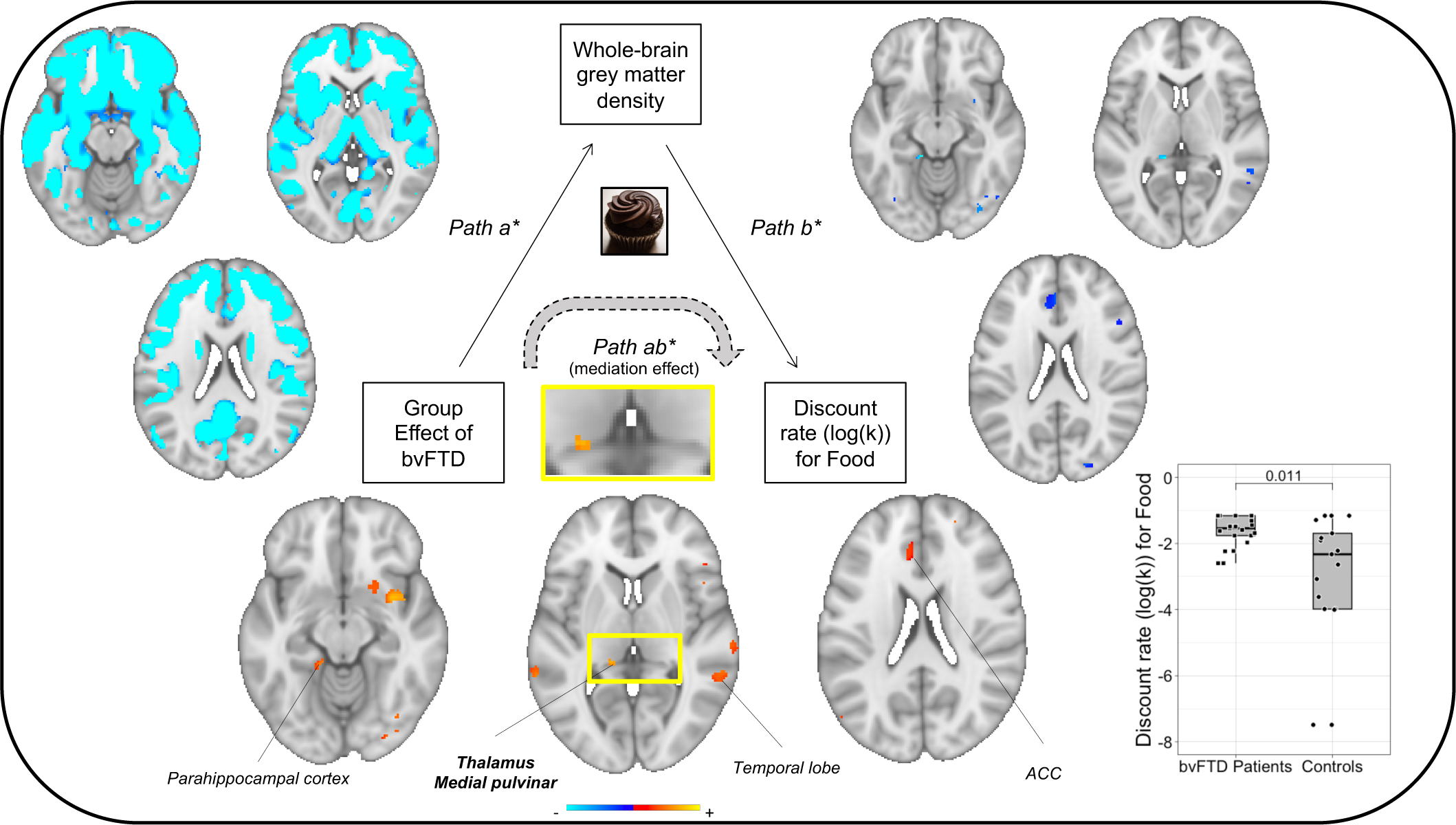
Neuroanatomical whole-brain mediators for the effect of bvFTD on discounting of food rewards. The path diagram displays the voxels of grey matter density (GMD) that significantly contributed to each path of the mediation model. The blue voxels in *path a* reflect the decreased GMD in bvFTD (n=21) compared to healthy controls (n=17), which corresponds to the typical atrophy pattern observed in bvFTD. *Path b* shows the voxels in which GMD is correlated to the discount rates across all participants (n=38) by controlling for the effect of group. Voxels in blue correspond to a negative correlation in which lower grey matter density contributes to higher discounting. *Path ab* represents brain regions in which GMD mediates the effect of bvFTD on discount rates. Voxels in yellow correspond to positive mediators that positively contribute to the increase in discounting due to bvFTD. *All results are FDR-corrected and thresholded at q<0.05 across the three paths. The graph on the left shows the significant direct effect of bvFTD condition (compared to healthy controls) on the discount rate of food rewards.

We were particularly interested in clusters significant across the three paths for both money and food rewards. Among these clusters, some were detected in the same locations for both rewards: in the left medial pulvinar thalamic nucleus, in the left parahippocampal cortex, and in the right temporal cortex. In these three regions, bvFTD patients presented significant grey matter atrophy (path a), lower GMD was associated with higher discount rate (path b), and the loss of GMD due to bvFTD positively contributed to increasing the discount rate (path ab). Of note, among clusters significant across the three paths, medial pulvinar was systematically associated with the largest effect size as a positive mediator of the effect of bvFTD on delay discounting (see Supplementary Tables 3 and 4 for path ab). For both reward types, correlations between the average grey matter density in the medial pulvinar cluster and the discount rates in the bvFTD and control groups are shown in Supplementary Fig. 6.

For food rewards, the mediation analysis also revealed two additional clusters significant across paths a, b and ab: one in the left orbitofrontal cortex (OFC) and one in the anterior cingulate cortex (ACC).

## DISCUSSION

Behavioural changes, including impulsivity symptoms, have been consistently reported in bvFTD, but the cognitive and neural mechanisms underlying these symptoms are still debated. In this paper, we aimed to advance our understanding of bvFTD-related behavioural changes by focusing on a specific facet of impulsivity in decision-making: delay discounting, or the tendency to prefer smaller sooner over larger later rewards. We used data from two intertemporal choice tasks (one with monetary stimuli and one with food stimuli) in bvFTD patients and healthy controls to assess their discount rates (i.e., their degree of impulsivity). For both money and food rewards, we found increased discount rates in bvFTD patients compared to healthy controls. BvFTD symptoms of disinhibition and eating behaviour changes were related to higher discount rates, thus confirming the assumed link between these core behavioural symptoms and a consistent preference for smaller sooner rewards. Moreover, atrophy and reduced grey matter density in the thalamus (medial pulvinar nucleus) and other regions of the limbic system (parahippocampal cortex, ACC and OFC) were found to play a key role in the alteration of the discount rate. Better understanding of the behavioural changes and neural structures associated with increased discounting in bvFTD might provide clinical insights into the general mechanisms of impulsivity across conditions. By uncovering mechanisms of impulsive decision-making, these findings could also indirectly contribute to clinical applications, especially for the treatment of the neuropsychological mechanisms potentially underlying the behavioural symptoms in bvFTD.

Discount rates were higher in bvFTD patients than in controls for both money and food rewards. This confirmed previous findings in bvFTD^13–17,32^ and also allowed us to verify that increased delay discounting generalizes across different rewards in bvFTD. This was found to be connected with a lack of sensitivity to the value of the larger later options, in accordance with Frost and McNaughton’s theory of delay discounting.^33^ Indeed, steeper discounters are assumed to be those for whom the delayed rewards are less salient (and therefore less likely to be chosen): for these individuals, reward salience intensely decreases with the perceived mental distance to reach it.^33^

BvFTD patients’ behavioural symptoms potentially indicative of an impulsive preference for immediacy were related to their higher impatience for monetary and food rewards in a decision-making task. First, higher discount rates were linked to higher deficits of inhibition, indicative of a preference for the immediate prepotent answers. Second, higher discount rates were associated with more eating behaviour changes, suggestive of higher preference for immediate food rewards. Therefore, the preference for immediacy of outcome might be a central common mechanism explaining different behavioural symptoms of bvFTD (including executive apathy, which may indicate preference for immediate answers to distractors instead of focusing on long-term goal management). The reduced attraction of delayed rewards associated with increased discount rates might be part of a more general mechanism of reduced attention to the most distant answers to any stimulation. By causing a decrease in attention towards the semantically distant words, this mechanism may explain patients’ difficulty in inhibiting the semantically close words in the Hayling test.^26^ Of note, individual differences in this type of inhibition deficit were linked to differences in discounting of money rewards among bvFTD patients. This finding is consistent with observations of a previous study that used factor analysis to identify impulsivity dimensions in patients with Parkinson’s disease: they identified a factor of preference for immediacy highly related to both discount rates of money and performance on the Hayling test.^34^

Delay discounting of food rewards was more closely associated with eating behaviour changes than with inhibition deficits. With food stimuli, the discount rate might not reflect only the general preference for more immediate reward outcomes; it might also reflect the specific changes in appetite regulation and sensitivity to food often observed in bvFTD patients.^35^ Results suggest that steeper discounting of food rewards is mostly associated with dimensions of lack of inhibition in eating behaviour, in particular the EBI ‘food approach’ subscale^29^ or the tendency to rush on food (which tends to be positively associated with discounting of food rewards among bvFTD patients). Changes in food preferences might be explained less by changes in impulsivity (captured by a steeper discount rate) and more related to compulsive stereotypical patterns of behaviour.

Grey matter density in a specific nucleus of the thalamus, the medial pulvinar, was related to discount rates (of both money and food) after controlling for group effect (path b), and also evidenced as an important mediator of the increase in discount rate due to bvFTD (path ab). Frost and McNaughton proposed that the ‘crude’ positive subjective value of the delayed gain is linked to activity in the thalamus, a key node in feedback loops between basal ganglia, hippocampus, and frontal and prefrontal cortex.^33^ Results of the whole-brain mediation did not evidence the OFC as the most prominent and consistent mediator of altered discounting, contrary to what we expected. However, thalamic lesions must impact some of the OFC–basal ganglia loops, which are associated with diverse neuropsychiatric symptoms including impulsive and compulsive disorders.^36,37^

The OFC structure was related to the discount rate independent of group effect and explained bvFTD patients’ increased discounting only with food rewards. The OFC has been proposed as a neural substrate that encodes value from a set of attributes attached to a particular object.^38–43^ It is part of a valuation network involved in delay discounting through its role in encoding the subjective values of both the immediate and delayed rewards.^44,45^ We found many regions related to reward processing (such as the OFC) among mediators of the increased discount rates of food in bvFTD. It is likely that differences in discounting of food are driven partly by bvFTD-related changes in sensitivity to the specific food rewards (associated with the symptom of eating behaviour changes) and not only by changes in value and salience of immediate versus delayed options. Thus, in bvFTD, the involvement of reward processing networks in the resulting discount rate might be particularly important with food rewards. Interestingly, among regions that positively mediated the effect of bvFTD on discounting only with food, the right OFC, ventral insula and striatum correspond to areas previously involved in binge eating in bvFTD patients.^46^ Some regions found as positive mediators only with food (ACC, insula) also overlap with the most important contributors of a brain marker predicting food and drug cravings.^47^

Among regions in which atrophy was found to alter delay discounting, both the medial pulvinar and the ACC are involved in both the connectivity of the salience network and the processing of conflict. BvFTD-associated atrophy maps the salience network, an intrinsic connectivity network that functions to represent the emotional significance of internal and external stimuli in order to guide behavioural responses.^48–50^ The medial pulvinar is highly connected with salience network nodes such as the anterior insula and ACC,^51^ and damage to the pulvinar has been shown to disrupt functional connectivity in the salience network in bvFTD patients.^52^ Atrophy to the medial pulvinar and ACC may thus increase impulsive decision-making by causing dysfunctions of the salience network, thereby reducing the positive salience of larger later rewards.

Moreover, the medial pulvinar is assumed to coordinate the processing of concurrent information and play a key role in determining decisions in conflictual situations with competing constraints, as in the delay discounting task.^53,54^ Similarly, the ACC with its connectivity to other brain regions, such as the lateral prefrontal and supplementary motor areas, has been related to conflict processing.^55^ Thus, evidencing these two regions as mediators of the alteration of delay discounting in bvFTD is consistent with the assumption that the impairment of conflict detection is a central mechanism underlying impulsive choices in this condition.^12^

Apart from the medial pulvinar, the OFC and the ACC, the parahippocampal cortex was also related to delay discounting (of both monetary and food rewards) independent of group effect and mediated the alteration of discounting in bvFTD. The parahippocampal cortex is involved in the consolidation of episodic memory, known to impact delay discounting.^56^ Along with other regions of the medial temporal lobe, it is assumed to contribute to a prospection network that mediates individual differences in discount rate through its role in simulating future outcomes.^44^ Evidencing the parahippocampal cortex as a mediator suggests that the alteration of discounting in bvFTD is not due only to impairments of reward valuation and salience; it also involves deficits in the contextualization of potential rewards.

From a clinical point of view, future studies could investigate how to use delay discounting for the treatment and diagnosis of bvFTD. Delay discounting and its structural neural bases could constitute targets in the development of treatments for multiple core behavioural symptoms of bvFTD. For instance, novel treatments targeting abnormalities in the medial pulvinar through invasive and non-invasive brain stimulation might have indirect effects to reduce symptoms of impulsivity in early bvFTD as in some psychiatric conditions.^37^

Further, delay discounting could become an interesting marker for the early diagnosis of bvFTD^12^ by contributing to the monitoring of non-symptomatic individuals at risk of FTD due to genetic mutations. The medial pulvinar has a significant role in the pathogenesis of bvFTD, especially in carriers of the C9orf72 genetic mutation, who may present medial pulvinar atrophy due to a developmental lesion.^52,57^ Abnormal development of the medial pulvinar also constitutes a common factor of several neurodevelopmental diseases involving impulsivity disorders^53,54^ that may predispose to developing bvFTD.^12^ Moreover, subtle behavioural changes in inhibition deficits appear very early in non-symptomatic individuals carrying a C9orf72 mutation predisposing to FTD.^58^ Since delay discounting is sensitive to both medial pulvinar lesions and inhibition deficits, it could be investigated as a complementary tool providing added value for the early diagnosis of bvFTD.

A few methodological limitations of our study, however, imply the need for further investigation to provide more evidence for our results and their implications. The main limitation of this study was the necessity to approximate several discounting values in cases of unique choice patterns (either only SS or only LL options) and cases of very inconsistent choices especially in bvFTD patients. It is possible that our paradigm was not adapted enough to reflect a very large range of discounting parameters, in particular very high discount rates in bvFTD. The actual range of discount rates might be larger in bvFTD patients. Of note, even without these approximations, delay discounting parameters were found higher in bvFTD patients than in controls (significantly for food). Moreover, the significant relationship between discount rates and lower sensitivity to LL reward further confirmed the validity of our approximations. Another methodological limitation of the study was related to the use of subjective reports to measure eating behaviour changes. Lack of variance in measures of delay discounting in bvFTD (in particular for the food paradigm) as well as subjective biases in measures of eating behaviour changes may have prevented us from evidencing the sensitivity of delay discounting to individual differences of eating behaviour changes within the group of bvFTD patients. Future studies would therefore benefit from the use of adjusted paradigms of delay discounting and more objective assessments of eating behaviour changes in bvFTD.

In conclusion, the results of this study have implications for a better understanding of impulsivity in bvFTD, which might extend to other conditions. The preference for smaller sooner rewards measured by discount rates may relate to a common factor of preference for immediacy underlying core bvFTD symptoms of impulsivity, including in the domain of eating. Atrophy to the medial pulvinar, by disrupting saliency detection of the delayed distant options, may causally contribute to the strong preference for immediacy in bvFTD. This study questions the central role of the OFC, which has often been suggested to explain impulsivity in bvFTD, and opens a path to further investigation of other brain regions, such as the medial pulvinar and its mediating role across neurodegenerative diseases and psychiatric conditions. For instance, do differential levels of medial pulvinar atrophy explain the differences in impatience for reward and impulsivity observed in Alzheimer’s disease and in bvFTD?^12^ In terms of clinical applications, medial pulvinar atrophy could be investigated as a transdiagnostic marker of impulsivity and as a potential therapeutic target for the treatment of impulsivity troubles across neurodegenerative and psychiatric conditions.

## Data availability

Data and analysis scripts used for the delay discounting tasks are available on OSF (https://osf.io/embr3/). Structural MRI data and other clinical data are available from the corresponding author, upon reasonable request.

## Abbreviations

ACC: anterior cingulate cortex AD = Alzheimer’s disease
bvFTD: behavioural variant frontotemporal dementia EBI = Eating Behaviour Inventory
GMD: grey matter density
HSCT: Hayling Sentence Completion Test ITC = intertemporal choice
LL: larger later
MMSE: Mini-Mental State Evaluation OFC = orbitofrontal cortex
PFC: prefrontal cortex SS = smaller sooner
VBM: voxel-based morphometry vmPFC = ventromedial prefrontal cortex

## Acknowledgements

We would like to thank patients, caregivers and organizers (in particular, Armelle Rametti-Lacroux) of the ECOCAPTURE consortium study and all the students who contributed to help data collection.

## Supplementary material

Supplementary material is available at Brain online.

## Author contributions

Study conception and design: HP, LS, BB, RL. Data acquisition: BB, AD. Analysis and interpretation of data: VG, AD under supervision of HP, LK, LS. Study supervision: HP, VG. Writing the first version of the manuscript: VG, HP. Obtaining funding: HP, RL, BB. All authors critically revised the manuscript for its intellectual content and approved its final version.

## Ethics approval

This study is part of clinical trial C16-87 sponsored by INSERM, the French national institute for biomedical research. It was granted approval by the local Ethics Committee (‘Comité de Protection des Personnes’) on 17 May 2017 and registered in a public registry (Clinicaltrials.gov: NCT03272230).

## Consent to participate

All study participants gave their written informed consent to participate, according to the Declaration of Helsinki and in line with French ethical guidelines.

