## Supplementary for "High delay discounting relates to core symptoms and to pulvinar atrophy in frontotemporal dementia"

### SUPPLEMENTARY MATERIALS

**Additional Methods: Sensitivities to larger later reward for money and food**

For each participant and each reward type, we computed the individual sensitivity to larger later (LL) reward as a complementary metric, useful in particular for the validation of computed discount rates. For this purpose, we fitted a logistic regression model in each participant predicting the trial-to-trial probability of choosing the LL option from the LL reward amount and LL delay value. We used the regression coefficient of LL reward amount as an estimate of the individual sensitivity to LL reward (higher sensitivity to reward corresponding to higher values).

Using nonparametric Spearman rank correlations, we tested the links between the computed discount rates and sensitivity to LL reward (including after correction for group effect) to verify that higher discount rates were associated with lower sensitivity to LL reward. Further, if bvFTD patients are indeed more impatient for reward regardless of the amount offered by the LL option, they should also show lower sensitivity to LL reward. We used nonparametric Wilcoxon tests to compare bvFTD patients and controls on sensitivity to LL reward calculated for money and food rewards.

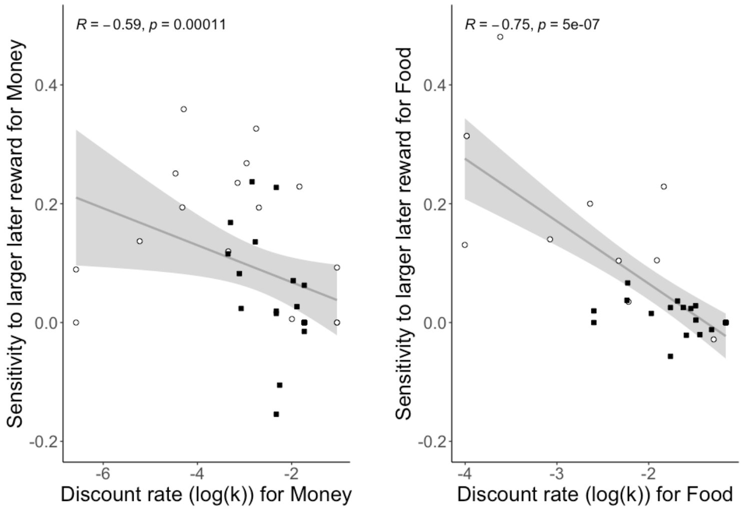

**Supplementary Figure 1. Correlations between discount rates (log(k)) and sensitivities to larger later reward for money and food.**

On the left: Spearman correlation between the discount rate for Money and the sensitivity to larger later reward for Money across bvFTD patients (N=22; represented as black squares) and controls (N=16; represented as white circles). On the right: Spearman correlation between the discount rate for Food and the sensitivity to larger later reward for Food across bvFTD patients (N=20 after removing one extreme outlier on sensitivity to larger later reward; represented as black squares) and controls (N=13 after removing two extreme outliers on sensitivity to larger later reward and two extreme outliers on discount rate; represented as white circles).

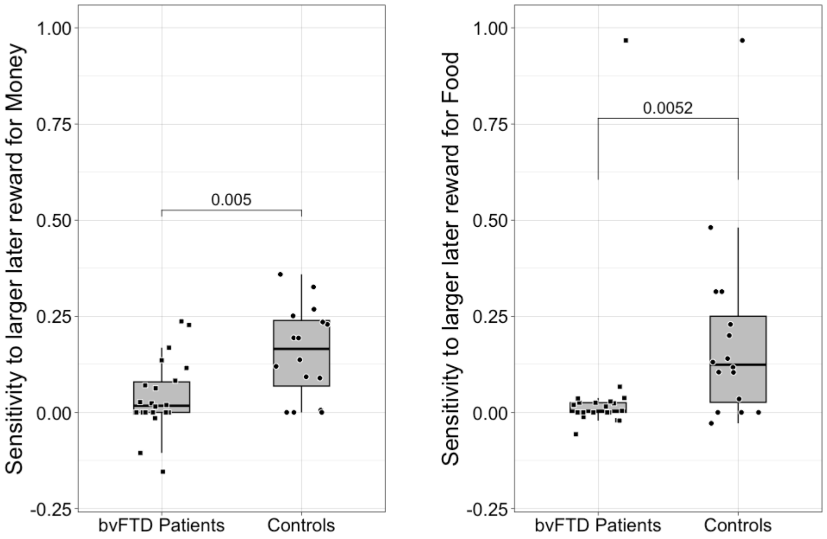

**Supplementary Figure 2. Effect of bvFTD on sensitivity to larger later reward.**
On the left: Wilcoxon test of the difference between bvFTD patients (N=22) and controls (N=16) on sensitivity to larger later reward for Money; sensitivity to reward was lower in bvFTD patients than in controls. On the right: Wilcoxon test of the difference between bvFTD patients (N=21) and controls (N=17) on sensitivity to larger later reward for Food; sensitivity to reward was lower in bvFTD patients than in controls. NB: for display purposes, one extreme outlier on sensitivity to larger later reward for Food was not represented on the graph, and the *p*-value displayed on the graph corresponds to the obtained value after removing this extreme outlier.

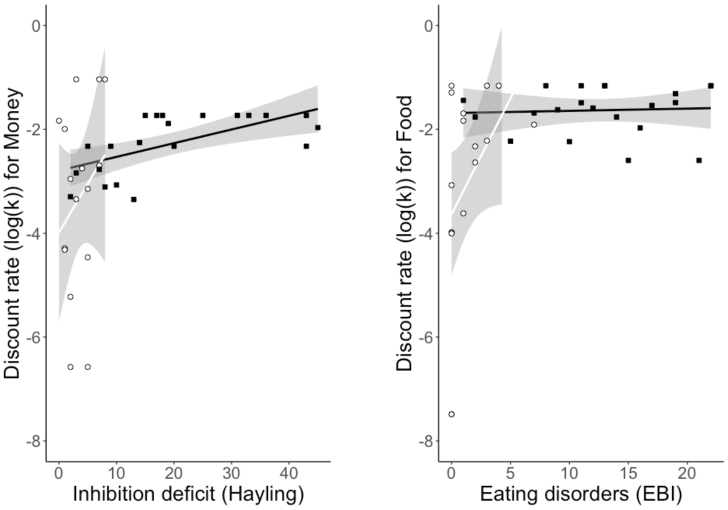

**Supplementary Figure 3. Correlations between discount rates and impulsivity-related symptoms within each group.**

On the left: Spearman correlations between inhibition deficit and discount rate for Money by participant group, i.e., among bvFTD patients (N=22; represented as black squares) and among controls (N=16; represented as white circles); discount rate for Money is sensitive to individual differences in inhibition deficit among bvFTD patients (R=0.67, p=0.0009) but not among controls (R=0.22, p=0.42). On the right: Spearman correlations between eating behaviour changes and discount rate for Food by participant group, i.e., among bvFTD patients (N=21; represented as black squares) and among controls (N=17; represented as white circles); discount rate for Food is sensitive to individual differences in eating behaviour changes among controls (R=0.52, p=0.03) but not among bvFTD patients (R=0.15, p=0.51).

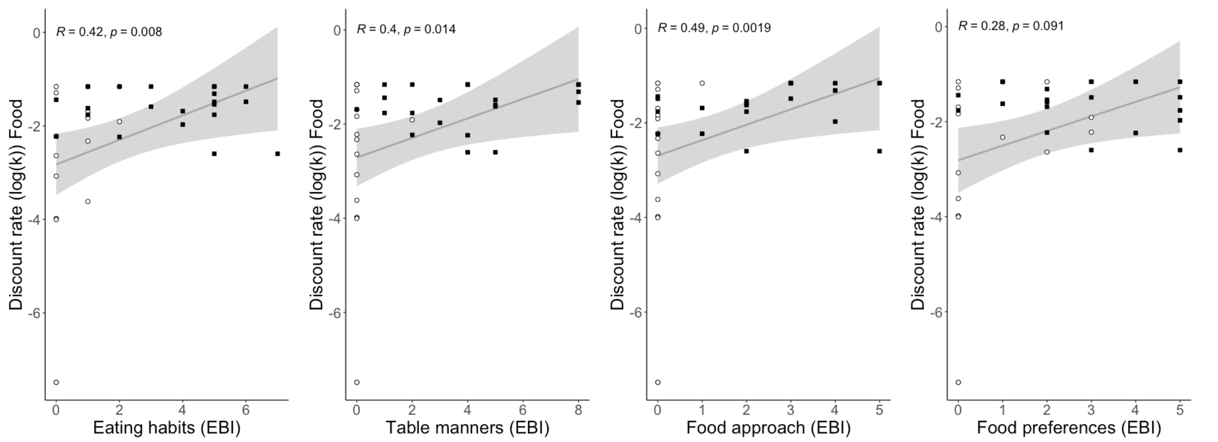

**Supplementary Figure 4. Correlations between discount rate for food and the four subscales of the Eating Behavior Inventory.**

From left to right: Spearman correlation between discount rate for Food and changes in eating habits across bvFTD patients (N=21; represented as black squares) and controls (N=17; represented as white circles); Spearman correlation between discount rate for Food and changes in table manners across bvFTD patients (N=21; represented as black squares) and controls (N=17; represented as white circles); Spearman correlation between discount rate for Food and changes in food approach tendency across bvFTD patients (N=21; represented as black squares) and controls (N=17; represented as white circles); Spearman correlation between discount rate for Food and changes in food preferences across bvFTD patients (N=21; represented as black squares) and controls (N=17; represented as white circles).

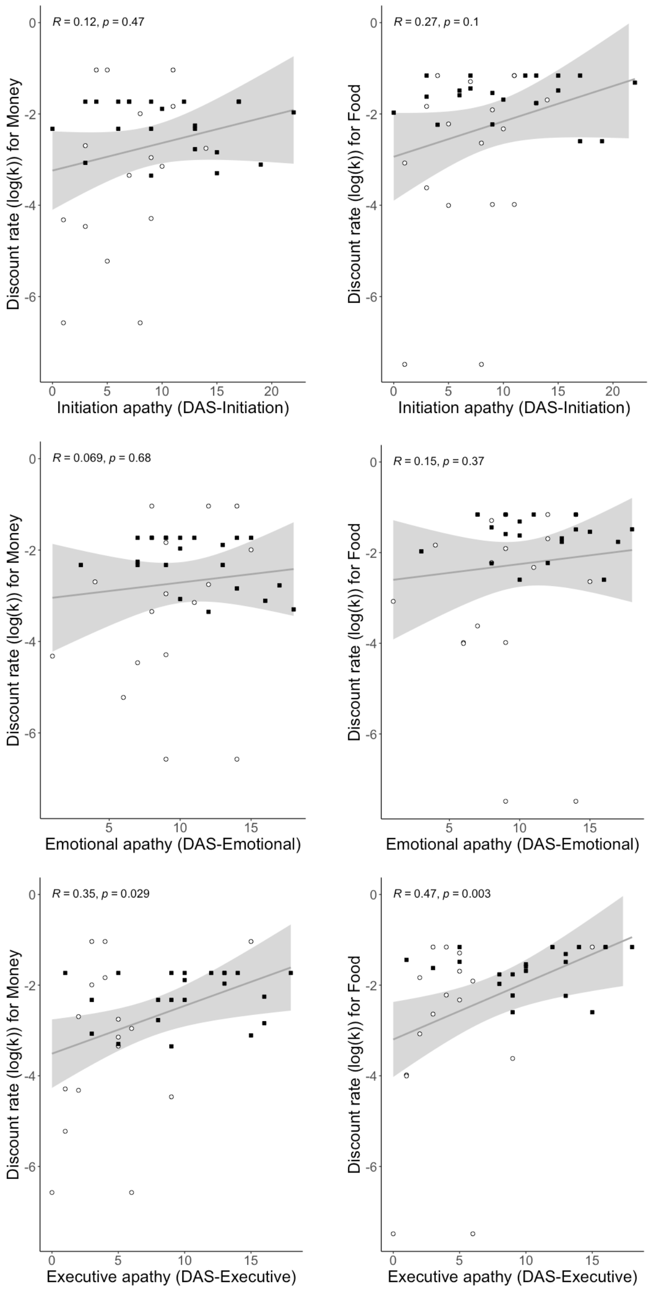

**Supplementary Figure 5. Correlations between discount rates and apathy dimensions of the Dimensional Apathy Scale (DAS).**

From top left to bottom right: Spearman correlation between discount rate for Money and initiation apathy across bvFTD patients (N=22; represented as black squares) and controls (N=16; represented as white circles); Spearman correlation between discount rate for Food and initiation apathy across bvFTD patients (N=21; represented as black squares) and controls (N=17; represented as white circles); Spearman correlation between discount rate for Money and emotional apathy across bvFTD patients (N=22; represented as black squares) and controls (N=16; represented as white circles); Spearman correlation between discount rate for Food and emotional apathy across bvFTD patients (N=21; represented as black squares) and controls (N=17; represented as white circles); Spearman correlation between discount rate for Money and executive apathy across bvFTD patients (N=22; represented as black squares) and controls (N=16; represented as white circles); Spearman correlation between discount rate for Food and executive apathy across bvFTD patients (N=21; represented as black squares) and controls (N=17; represented as white circles).

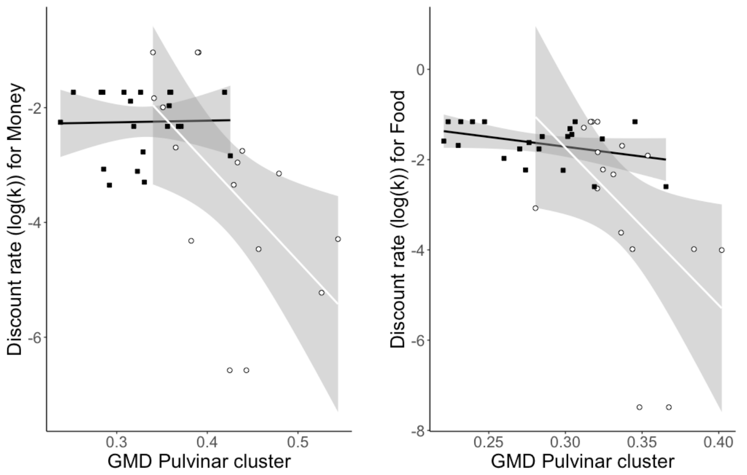

**Supplementary Figure 6. Correlations between average grey matter density (GMD) in the medial pulvinar cluster and discount rate within each group.**
On the left: Spearman correlations between GMD in medial pulvinar cluster and discount rate for Money by participant group, i.e., among bvFTD patients (N=22; represented as black squares) and among controls (N=16; represented as white circles); differences in GMD in the medial pulvinar are significantly related to individual differences of discount rate for Money among controls (R=-0.64, p=0.008) but not among bvFTD patients (R=-0.05, p=0.83). On the right: Spearman correlations between GMD in the medial pulvinar cluster and discount rate for Food by participant group, i.e., among bvFTD patients (N=21; represented as black squares) and among controls (N=17; represented as white circles); differences in GMD in the medial pulvinar are significantly related to individual differences in discount rate for Food among controls (R=-0.67, p=0.003) but not among bvFTD patients (R=-0.19, p=0.42).

**Supplementary Table 1. Demographic and main clinical measures of bvFTD patients and controls.**

Data are given as mean (SD). BvFTD patients: N= 22/controls: N= 17. For comparison, we used Wilcoxon tests for non-normally distributed variables and Student's t-tests for normally distributed variables. First, the main clinical measures are listed. MMSE: Mini-Mental State Examination; DRS: Dementia Rating Scale; FAB: Frontal Assessment Battery. Second, variables of interest in the study are presented. Hayling – error: objective measure of inhibition deficit from the Hayling Sentence Completion Test (number of errors in the inhibition phase of the test); EBI: Eating Behavior Inventory, global measure of changes in eating behaviour. DAS: Dimensional Apathy Scale.

| | bvFTD | Controls | bvFTD vs. controls |
|---|---|---|---|
| % women | 36.4% | 52.9% | $\chi^2 = 0.5$ ; $p = 0.48$ |
| Age | 66.5 (8.5) | 62.2 (7.2) | W = 256 ; p = 0.05 |
| Education level | 6.1 (2.0) | 7.2 (1.1) | W = 136 ; p = 0.13 |
| MMSE (/30) | 23.8 (2.6) | 29.5 (0.7) | W = 5.5 ; p < 0.001 |
| DRS (/144) | 119.4 (8.9) | 142.2 (1.3) | W = 0 ; p < 0.001 |
| FAB (/18) | 12.1 (3.4) | 17.4 (0.9) | W = 6.5 ; p < 0.001 |
| Hayling – error | 19.8 (13.7) | 3.3 (2.5) | W = 331.5 ; p < 0.001 |
| EBI (/32) | 13.1 (6.2) | 1.4 (1.9) | W = 360 ; p < 0.001 |
| EBI – Eating habits (/8) | 3.7 (2.1) | 0.5 (0.7) | W = 334.5 ; p < 0.001 |
| EBI – Food preferences (/8) | 2.9 (1.7) | 0.7 (1.1) | W = 317 ; p < 0.001 |
| EBI – Table manners (/8) | 4.0 (2.5) | 0.1 (0.5) | W = 360.5 ; p < 0.001 |
| EBI – Food approach (/8) | 2.5 (1.7) | 0.1 (0.2) | W = 337 ; p < 0.001 |
| DAS-Initiation (/24) | 10.4 (5.7) | 7.1 (3.8) | T = 2.2 ; p = 0.04 |
| DAS-Emotional (/24) | 10.9 (3.8) | 9.1 (3.7) | T = 1.5 ; p = 0.1 |
| DAS-Executive (/24) | 10.0 (4.6) | 4.2 (3.6) | W = 311.5 ; p < 0.001 |

**Supplementary Table 2. Combination of SS, LL amounts, and delays used in the two delay discounting paradigms with money and food stimuli.**
The presentation order was randomized for each participant. Indifference k denotes the discounting rate at which the SS and LL options should be chosen at equal proportions.

| SS delay (in days) | LL delay (in days) | SS amount (in euros or chocolates) | LL amount (in euros or chocolates) | Indifference k |
|---|---|---|---|---|
| 0 | 14 | 8 | 10 | 0.017857143 |
| 0 | 14 | 8 | 32 | 0.214285714 |
| 0 | 14 | 17.5 | 20.5 | 0.012244898 |
| 0 | 14 | 21 | 21.25 | 0.00085034 |
| 0 | 14 | 22.5 | 32.5 | 0.031746032 |
| 0 | 14 | 23 | 25.75 | 0.008540373 |
| 0 | 14 | 23.5 | 23.7 | 0.000607903 |
| 0 | 14 | 27 | 36 | 0.023809524 |
| 0 | 14 | 28 | 56 | 0.071428571 |
| 0 | 28 | 12 | 13.5 | 0.004464286 |
| 0 | 28 | 12 | 24 | 0.035714286 |
| 0 | 28 | 14 | 14.2 | 0.000510204 |
| 0 | 28 | 15 | 20 | 0.011904762 |
| 0 | 28 | 16 | 23 | 0.015625 |
| 0 | 28 | 17.1 | 17.2 | 0.000208855 |
| 0 | 28 | 21 | 24.5 | 0.005952381 |
| 0 | 28 | 24 | 96 | 0.107142857 |
| 0 | 28 | 31 | 39 | 0.00921659 |
| 14 | 28 | 10.15 | 10.3 | 0.001071429 |
| 14 | 28 | 15.75 | 17.5 | 0.008928571 |
| 14 | 28 | 17 | 24 | 0.05 |
| 14 | 28 | 18 | 24 | 0.035714286 |
| 14 | 28 | 21.1 | 21.2 | 0.000340136 |
| 14 | 28 | 28 | 33 | 0.01552795 |
| 14 | 28 | 35 | 44 | 0.024725275 |
| 28 | 42 | 9 | 10.5 | 0.017857143 |
| 28 | 42 | 11 | 13.75 | 0.035714286 |
| 28 | 42 | 14.05 | 14.1 | 0.000256016 |
| 28 | 42 | 22.25 | 22.65 | 0.001332001 |
| 28 | 42 | 24 | 32 | 0.071428571 |
| 28 | 42 | 26 | 29 | 0.010714286 |
| 28 | 42 | 28 | 40 | 0.214285714 |

**Supplementary Table 3. Detailed results of whole-brain mediation analyses with money rewards.**

For paths b and ab, we list all the clusters significant at q<0.05 (= p<0.008) FDR-corrected (across paths a, b and ab) corresponding to both positive and negative effects. Atlas label: reference region with highest number of in-region voxels. Volume: volume of contiguous region in cubic mm. X, Y and Z: peak coordinates in MNI space. Max(z): signed max over p. **Clusters in bold correspond to cortical and subcortical regions of overlap between paths a, b and ab.**

***Whole-brain mediation: Money – Path b***

*Positive effects*

| Name | Atlas label | Volume | X | Y | Z | max(z) |
|---|---|---|---|---|---|---|
| Precuneous cortex | Ctx_31pd_R | 56 | 12 | -53 | 32 | 7.0345 |
| Inferior frontal gyrus | Ctx_45_L | 80 | -53 | 27 | -2 | 7.0345 |
| Lateral occipital cortex | Ctx_MIP_L | 504 | -18 | -66 | 48 | 7.0345 |
| Supramarginal gyrus | Ctx_PFt_R | 1016 | 53 | -36 | 51 | 7.0345 |
| Precuneous cortex | Ctx_7Am_R | 264 | 8 | -57 | 65 | 7.0345 |
| Precuneous cortex | Ctx_POS2_R | 848 | 5 | -72 | 44 | 7.0345 |
| Ventromedial prefrontal cortex | Ctx_10v_L | 136 | -3 | 27 | -24 | 7.0345 |
| Precentral gyrus | Ctx_4_L | 376 | -6 | -30 | 53 | 7.0345 |
| Precuneous cortex | Ctx_23c_L | 976 | -14 | -36 | 42 | 7.0345 |
| Occipital fusiform gyrus | Ctx_V4_R | 256 | 26 | -78 | -6 | 7.0345 |
| Occipital pole | Ctx_V1_R | 392 | 21 | -98 | -2 | 6.7649 |
| Precuneous cortex | Ctx_7Pm_R | 64 | 2 | -63 | 53 | 6.7551 |
| Middle frontal gyrus | Ctx_8C_L | 40 | -38 | 21 | 30 | 6.4902 |
| Angular gyrus | Ctx_7PC_L | 112 | -45 | -54 | 53 | 6.0388 |
| Precuneous cortex | Ctx_PCV_L | 96 | -8 | -53 | 53 | 4.0588 |
| Occipital pole | Ctx_V3_L | 96 | -26 | -95 | 3 | 3.6701 |

*Negative effects*

| Name | Atlas label | Volume | X | Y | Z | max(z) |
|---|---|---|---|---|---|---|
| Cerebellum | Cblm_VI_L | 976 | -29 | -39 | -33 | -22.125 |
| Cerebellum | Cblm_VIIIa_R | 4936 | 11 | -72 | -45 | -17.87 |
| Cerebellum | Cblm_Vermis_VI | 576 | -2 | -74 | -14 | -15.356 |
| Cerebellum | Cblm_I_IV_L | 64 | -11 | -44 | -5 | -14.873 |
| **Thalamus pulvinar** | **Thal_Pulv** | **256** | **-14** | **-35** | **3** | **-14.581** |
| **Parahippocampal cortex** | **Ctx_PHA1_L** | **168** | **-17** | **-38** | **-14** | **-13.811** |
| **Middle temporal gyrus** | **Ctx_STV_R** | **240** | **65** | **-47** | **11** | **-12.806** |
| Inferior temporal gyrus | Ctx_TE2p_L | 32 | -50 | -42 | -15 | -12.562 |

| | | | | | | |
|---|---|---|---|---|---|---|
| Cerebellum | Cblm_VIIIa_L | 704 | -12 | -69 | -51 | -10.549 |
| Cerebellum | Cblm_CrusI_L | 984 | -24 | -83 | -27 | -8.4412 |
| Cerebellum | Cblm_IX_R | 48 | 8 | -53 | -35 | -7.71 |
| Angular gyrus | Ctx_IP2_R | 272 | 41 | -50 | 47 | -7.192 |
| Cerebellum | Cblm_CrusII_L | 464 | -33 | -72 | -42 | -6.4424 |
| Occipital pole | Ctx_V1_L | 96 | -11 | -93 | 11 | -6.2466 |
| Lateral occipital cortex | Ctx_FST_R | 104 | 45 | -69 | -6 | -6.1674 |
| Lateral occipital cortex | Ctx_MT_L | 48 | -44 | -72 | 9 | -5.2595 |
| Occipital fusiform gyrus | Ctx_V3_L | 88 | -18 | -71 | -9 | -5.2073 |
| **Occipital fusiform gyrus** | **Ctx_V8_L** | **112** | **-35** | **-71** | **-14** | **-4.0465** |
| Angular gyrus | Ctx_PGi_R | 488 | 50 | -56 | 26 | -3.6751 |

***Whole-brain mediation: Money – Path ab***

*Positive effects*

| Name | Atlas label | Volume | X | Y | Z | max(z) |
|---|---|---|---|---|---|---|
| **Thalamus pulvinar** | **Thal_Pulv** | **296** | **-14** | **-33** | **3** | **0.98686** |
| Superior temporal gyrus | Ctx_STSdp_L | 192 | -56 | -23 | -3 | 0.89737 |
| Middle temporal gyrus | Ctx_TPOJ2_L | 384 | -48 | -57 | 5 | 0.85809 |
| Middle frontal gyrus | Ctx_9_46d_L | 424 | -23 | 36 | 29 | 0.75078 |
| Middle temporal gyrus | Ctx_TPOJ1_L | 104 | -66 | -42 | 3 | 0.7085 |
| Lateral occipital cortex | Ctx_TPOJ2_L | 80 | -51 | -66 | 9 | 0.69456 |
| **Middle temporal gyrus** | **Ctx_STV_R** | **672** | **60** | **-45** | **8** | **0.68241** |
| Parahippocampal gyrus | No label | 80 | -15 | 2 | -29 | 0.65357 |
| Precentral gyrus | Ctx_6a_R | 152 | 26 | -11 | 54 | 0.64242 |
| **Parahippocampal gyrus** | **Ctx_PHA1_L** | **384** | **-15** | **-38** | **-12** | **0.61398** |
| Cerebellum | Cblm_CrusI_L | 160 | -24 | -83 | -24 | 0.59469 |
| Cerebellum | Cblm_Vermis_VI | 272 | -3 | -74 | -9 | 0.55765 |
| Inferior temporal gyrus | Ctx_TE2p_L | 80 | -48 | -41 | -15 | 0.55005 |
| Insular cortex | Ctx_MI_R | 88 | 44 | 15 | -3 | 0.51757 |
| Cerebellum | Cblm_Vermis_VI | 40 | 2 | -72 | -12 | 0.47083 |
| Lateral occipital cortex | Ctx_V3CD_R | 64 | 30 | -78 | 12 | 0.46013 |
| Cerebellum | Cblm_CrusI_L | 112 | -20 | -81 | -33 | 0.42299 |
| **Occipital fusiform gyrus** | **Ctx_V8_L** | **64** | **-33** | **-69** | **-14** | **0.38635** |

*Negative effects*

| Name | Atlas label | Volume | X | Y | Z | max(z) |
|---|---|---|---|---|---|---|

| | | | | | | |
|---|---|---|---|---|---|---|
| Brain stem | Ctx_23c_L | 456 | -12 | -36 | 42 | -0.66327 |
| Lateral occipital cortex | Ctx_MIP_L | 264 | -18 | -66 | 50 | -0.48952 |
| Parietal operculum cortex | Ctx_PFcm_R | 280 | 44 | -33 | 23 | -0.29149 |

**Supplementary Table 4. Detailed results of whole-brain mediation analyses with food rewards.**

For paths b and ab, we list all the clusters significant at q<0.05 (= p<0.008) FDR-corrected (across paths a, b and ab) corresponding to both positive and negative effects. Atlas label: reference region with highest number of in-region voxels. Volume: volume of contiguous region in cubic mm. X, Y and Z: peak coordinates in MNI space. Max(z): signed max over p. **Clusters underlined in bold correspond to cortical and subcortical regions of overlap between paths a, b and ab.**

***Whole-brain mediation: Food – Path b***

*Positive effects*

| Name | Atlas label | Volume | X | Y | Z | max(z) |
|---|---|---|---|---|---|---|
| Superior frontal gyrus | Ctx_8Ad_L | 160 | -21 | 26 | 39 | 7.0345 |
| Superior frontal gyrus | Ctx_8Ad_R | 112 | 17 | 30 | 47 | 7.0345 |
| Lateral occipital cortex | Ctx_MIP_L | 112 | -18 | -63 | 48 | 7.0345 |
| Lateral occipital cortex | Ctx_FST_L | 40 | -48 | -66 | 0 | 7.0345 |
| Superior frontal gyrus | Ctx_6ma_L | 80 | -15 | 5 | 60 | 7.0345 |
| Precuneous cortex | Ctx_POS2_R | 304 | 3 | -72 | 42 | 7.0345 |
| Precuneous cortex | Ctx_7Pm_R | 96 | 0 | -62 | 53 | 7.0345 |
| Supramarginal gyrus | Ctx_PF_R | 208 | 54 | -38 | 51 | 7.0345 |
| Occipital fusiform gyrus | Ctx_V4_R | 336 | 26 | -78 | -8 | 7.0345 |
| Occipital pole | Ctx_V2_L | 296 | -6 | -89 | 15 | 7.0345 |
| Supramarginal gyrus | Ctx_AIP_R | 160 | 36 | -38 | 38 | 7.0297 |
| Cingulate gyrus | Ctx_23c_L | 96 | -12 | -36 | 42 | 6.8986 |
| Occipital pole | Ctx_V2_R | 168 | 23 | -96 | 2 | 6.1985 |
| Occipital pole | Ctx_V3_R | 280 | 12 | -89 | 17 | 6.1672 |
| Superior parietal lobule | Ctx_LIPv_L | 160 | -27 | -56 | 44 | 6.155 |
| Occipital fusiform gyrus | Ctx_V4_L | 64 | -26 | -74 | -5 | 5.9469 |
| Frontal pole | Ctx_9_46d_R | 200 | 27 | 47 | 11 | 5.418 |
| Superior frontal gyrus | Ctx_6ma_R | 80 | 17 | 2 | 63 | 4.7352 |
| Lateral occipital cortex | No label | 88 | -38 | -69 | 17 | 4.1177 |

*Negative effects*

| Name | Atlas label | Volume | X | Y | Z | max(z) |
|---|---|---|---|---|---|---|
| **Parahippocampal gyrus** | **Ctx_PHA1_L** | **160** | **-14** | **-38** | **-12** | **-23.802** |
| Cerebellum | Cblm_I_IV_L | 160 | -11 | -42 | -6 | -19.238 |
| Cerebellum | Cblm_CrusI_R | 1200 | 33 | -75 | -15 | -17.652 |
| Cerebellum | Cblm_VIIIa_R | 464 | 11 | -69 | -56 | -17.491 |
| Cerebellum | Cblm_I_IV_L | 200 | -2 | -44 | -9 | -16.741 |

| | | | | | | |
|---|---|---|---|---|---|---|
| **Thalamus pulvinar** | **Thal_Pulv** | **368** | **-15** | **-35** | **3** | **-15.909** |
| Cerebellum | Cblm_VIIIa_L | 360 | -12 | -69 | -56 | -13.109 |
| Cerebellum | Cblm_VIIIb_L | 1608 | -17 | -50 | -57 | -12.305 |
| Middle temporal gyrus | Ctx_STV_R | 96 | 66 | -44 | 9 | -11.996 |
| Precentral gyrus | Ctx_FEF_L | 472 | -50 | -9 | 47 | -11.21 |
| **Lateral occipital cortex** | **Ctx_TPOJ3_L** | **80** | **-53** | **-72** | **17** | **-10.464** |
| Lateral occipital cortex | Ctx_PH_R | 352 | 48 | -66 | -18 | -9.8397 |
| Middle temporal gyrus | Ctx_TE1p_L | 40 | -68 | -44 | 0 | -8.494 |
| Cerebellum | Cblm_CrusI_R | 216 | 26 | -86 | -27 | -8.3793 |
| Intracalcarine cortex | No label | 96 | 14 | -72 | 3 | -8.2769 |
| Superior parietal lobule | Ctx_AIP_R | 264 | 39 | -47 | 50 | -8.1443 |
| Middle temporal gyrus | Ctx_STSvp_R | 352 | 59 | -42 | -2 | -8.0273 |
| **Anterior cingulate gyrus** | **Ctx_p24_L** | **1048** | **-5** | **35** | **20** | **-7.8319** |
| **Middle temporal gyrus** | **Ctx_TPOJ1_R** | **376** | **57** | **-47** | **8** | **-7.7366** |
| Superior temporal gyrus | Ctx_A4_R | 248 | 68 | -26 | 3 | -7.7186 |
| Lateral occipital cortex | Ctx_PGp_R | 168 | 38 | -77 | 14 | -7.4899 |
| Cerebellum | Cblm_CrusII_R | 48 | 17 | -89 | -29 | -7.0886 |
| Cerebellum | Cblm_VI_R | 224 | 17 | -62 | -17 | -6.7382 |
| Lateral occipital cortex | Ctx_V3A_R | 504 | 23 | -87 | 24 | -6.7315 |
| Others | Ctx_Pir_R | 32 | 30 | 5 | -12 | -6.729 |
| Middle frontal gyrus | Ctx_55b_R | 48 | 51 | 6 | 48 | -6.3286 |
| Cerebellum | Cblm_V_R | 40 | 23 | -42 | -18 | -6.3265 |
| Cerebellum | Cblm_VIIIb_R | 120 | 26 | -41 | -51 | -6.2259 |
| Cerebellum | Cblm_VIIIb_R | 48 | 15 | -47 | -53 | -6.1876 |
| Occipital fusiform gyrus | Ctx_FFC_R | 112 | 39 | -68 | -11 | -6.0147 |
| **Frontal orbital cortex** | **Ctx_13l_L** | **128** | **-23** | **15** | **-27** | **-5.9295** |
| Cerebellum | Cblm_CrusI_L | 192 | -32 | -83 | -30 | -5.9118 |
| Temporal fusiform cortex | Ctx_PeEc_L | 248 | -26 | -8 | -45 | -5.6773 |
| Cerebellum | Cblm_CrusII_R | 856 | 12 | -83 | -36 | -5.4777 |
| Cerebellum | Cblm_CrusI_R | 112 | 39 | -62 | -20 | -5.358 |
| Occipital fusiform gyrus | Ctx_V3_R | 88 | 20 | -78 | -5 | -4.4783 |
| Middle temporal gyrus | Ctx_TPOJ2_R | 80 | 56 | -56 | 8 | -4.0816 |
| Temporal fusiform cortex | Ctx_TGv_R | 232 | 27 | -2 | -47 | -4.0095 |
| Occipital fusiform gyrus | Ctx_V8_L | 64 | -35 | -71 | -12 | -3.6114 |
| Inferior frontal gyrus | Ctx_IFJa_R | 272 | 48 | 20 | 23 | -3.4883 |
| Angular gyrus | Ctx_PGs_R | 64 | 50 | -57 | 30 | -2.6146 |

***Whole-brain mediation: Food – Path ab***

*Positive effects*

| Name | Atlas label | Volume | X | Y | Z | max(z) |
|---|---|---|---|---|---|---|
| Middle temporal gyrus | Ctx_STSdp_L | 1824 | -54 | -29 | -6 | 1.1806 |
| Cerebellum | Cblm_VIIIb_R | 792 | 17 | -45 | -47 | 1.0941 |
| Precentral gyrus | Ctx_4_L | 1584 | -53 | -9 | 45 | 1.0723 |
| Frontal orbital cortex | Ctx_13l_R | 3704 | 23 | 17 | -20 | 0.9856 |
| Planum polare | Ctx_TA2_L | 504 | -48 | -3 | -6 | 0.95925 |
| Putamen | Putamen_Pp_R | 2680 | 35 | 3 | -9 | 0.90532 |
| **Thalamus pulvinar** | **Thal_Pulv** | **352** | **-14** | **-38** | **3** | **0.90382** |
| Middle temporal gyrus | Ctx_FST_L | 144 | -48 | -56 | 3 | 0.88662 |
| Middle temporal gyrus | Ctx_TPOJ1_L | 472 | -65 | -42 | 5 | 0.86467 |
| Inferior frontal gyrus | Ctx_44_R | 216 | 48 | 17 | 11 | 0.85843 |
| **Middle temporal gyrus** | **Ctx_TPOJ1_R** | **1280** | **57** | **-42** | **3** | **0.80838** |
| Temporal fusiform cortex | Ctx_PeEc_L | 376 | -33 | -18 | -32 | 0.77288 |
| Angular gyrus | Ctx_STV_L | 472 | -62 | -51 | 12 | 0.75697 |
| Parahippocampal gyrus | Ctx_PeEc_R | 320 | 23 | 3 | -36 | 0.74826 |
| Middle temporal gyrus | Ctx_STSva_R | 96 | 51 | -14 | -20 | 0.71871 |
| **Frontal orbital cortex** | **Ctx_13l_L** | **1048** | **-18** | **18** | **-26** | **0.71837** |
| Temporal fusiform cortex | Ctx_TGv_R | 848 | 29 | -2 | -45 | 0.71542 |
| Frontal pole | Ctx_9_46d_R | 288 | 23 | 53 | 17 | 0.71522 |
| Putamen | Putamen_Pp_L | 616 | -32 | -2 | -6 | 0.70718 |
| Insular cortex | Ctx_MI_R | 280 | 42 | 15 | -3 | 0.69686 |
| Temporal pole | Ctx_TGd_R | 200 | 47 | 5 | -42 | 0.68051 |
| Parahippocampal gyrus | Ctx_PeEc_R | 1248 | 33 | -23 | -27 | 0.67486 |
| Middle frontal gyrus | Ctx_9_46d_L | 224 | -26 | 35 | 29 | 0.66799 |
| Precentral gyrus | Ctx_IFJp_L | 184 | -42 | 2 | 26 | 0.66603 |
| Cerebellum | Cblm_CrusI_L | 376 | -21 | -87 | -24 | 0.65883 |
| Cerebellum | Cblm_CrusI_R | 64 | 45 | -62 | -23 | 0.65186 |
| Middle frontal gyrus | Ctx_6a_R | 352 | 32 | -2 | 59 | 0.63603 |
| Cerebellum | Cblm_CrusI_R | 1056 | 36 | -71 | -18 | 0.6303 |
| **Parahippocampal gyrus** | **Ctx_PHA1_L** | **608** | **-17** | **-39** | **-12** | **0.62312** |
| Superior temporal gyrus | Ctx_A4_R | 584 | 66 | -23 | 9 | 0.62158 |
| Temporal pole | Ctx_TGv_L | 128 | -32 | 8 | -47 | 0.61643 |
| Lingual gyrus | Ctx_V2_L | 304 | -3 | -74 | -6 | 0.61559 |
| Precentral gyrus | Ctx_6mp_R | 64 | 21 | -11 | 59 | 0.6105 |
| Insular cortex | Ctx_MI_R | 40 | 41 | 5 | 0 | 0.6076 |
| Lateral occipital cortex | Ctx_PGi_L | 176 | -50 | -68 | 26 | 0.59054 |
| Middle frontal gyrus | Ctx_p9_46v_R | 320 | 47 | 24 | 26 | 0.51373 |
| Occipital fusiform gyrus | Ctx_V4_R | 256 | 27 | -86 | -15 | 0.51198 |
| Cuneal cortex | Ctx_V2_R | 32 | 3 | -71 | 26 | 0.51169 |
| Frontal pole | Ctx_8Ad_L | 152 | -24 | 36 | 42 | 0.50529 |
| Temporal pole | Ctx_TGv_L | 64 | -47 | 3 | -47 | 0.49609 |

| | | | | | | |
|---|---|---|---|---|---|---|
| Temporal fusiform cortex | Ctx_PeEc_L | 296 | -27 | -9 | -44 | 0.49153 |
| **Anterior cingulate gyrus** | **Ctx_p24_L** | **760** | **-6** | **32** | **23** | **0.48956** |
| Middle temporal gyrus | Ctx_A5_R | 216 | 68 | -27 | -2 | 0.48693 |
| Middle frontal gyrus | Ctx_8C_L | 136 | -45 | 24 | 41 | 0.48295 |
| **Lateral occipital cortex** | **Ctx_TPOJ3_L** | **264** | **-53** | **-72** | **18** | **0.48015** |
| Anterior cingulate gyrus | Ctx_p24_R | 144 | -2 | 32 | 17 | 0.47965 |
| Lateral occipital cortex | Ctx_PH_R | 112 | 48 | -63 | -18 | 0.47708 |
| Precentral gyrus | Ctx_4_L | 312 | -21 | -23 | 71 | 0.47068 |
| Cerebellum | Cblm_CrusI_L | 96 | -14 | -87 | -20 | 0.45977 |
| Temporal fusiform cortex | Ctx_FFC_L | 64 | -41 | -44 | -18 | 0.45682 |
| Middle temporal gyrus | Ctx_TE1a_R | 96 | 63 | -8 | -14 | 0.45362 |
| Frontal pole | Ctx_9a_R | 48 | 17 | 62 | 24 | 0.45017 |
| Frontal pole | Ctx_9_46d_R | 72 | 26 | 50 | 24 | 0.44951 |
| Putamen | Putamen_Pa_R | 136 | 23 | 14 | 3 | 0.44714 |
| Lingual gyrus | Ctx_V2_R | 144 | 5 | -77 | -8 | 0.44418 |
| Superior frontal gyrus | Ctx_9m_R | 120 | 5 | 47 | 33 | 0.41998 |
| Inferior frontal gyrus | Ctx_44_L | 128 | -56 | 12 | 17 | 0.41882 |
| Inferior frontal gyrus | Ctx_45_R | 136 | 48 | 29 | 5 | 0.40799 |
| Subcallosal cortex | Ctx_25_R | 24 | -2 | 20 | -11 | 0.40237 |
| Middle frontal gyrus | Ctx_8Av_R | 168 | 33 | 30 | 47 | 0.40166 |
| Caudate | Cau_R | 80 | 21 | 18 | 17 | 0.38119 |
| Middle frontal gyrus | Ctx_p9_46v_R | 112 | 39 | 35 | 36 | 0.37304 |
| Precentral gyrus | Ctx_6v_R | 472 | 59 | 2 | 38 | 0.3672 |
| Frontal orbital cortex | Ctx_47s_L | 272 | -33 | 23 | -21 | 0.34951 |
| Precentral gyrus | Ctx_6v_R | 64 | 60 | 5 | 27 | 0.30382 |
| Superior frontal gyrus | Ctx_8BM_L | 208 | -5 | 21 | 51 | 0.30002 |
| Middle frontal gyrus | Ctx_IFJa_L | 40 | -48 | 17 | 30 | 0.29962 |
| Superior parietal lobule | No label | 32 | 41 | -45 | 48 | 0.27238 |

*Negative effects*

| Name | Atlas label | Volume | X | Y | Z | max(z) |
|---|---|---|---|---|---|---|
| Frontal pole | Ctx_9_46d_R | 64 | 27 | 47 | 12 | -0.61739 |
| Cerebellum | Cblm_I_IV_L | 128 | -3 | -44 | -8 | -0.56971 |
| Supramarginal gyrus | Ctx_AIP_R | 176 | 35 | -38 | 39 | -0.41868 |
